# High definition DIC imaging uncovers transient stages of pathogen infection cycles on the surface of human adult stem cell-derived intestinal epithelium

**DOI:** 10.1101/2021.08.24.457471

**Authors:** Jorik M. van Rijn, Jens Eriksson, Jana Grüttner, Magnus Sundbom, Dominic-Luc Webb, Per M. Hellström, Staffan G. Svärd, Mikael E. Sellin

**Affiliations:** Science for Life Laboratory, Department of Medical Biochemistry and Microbiology, Uppsala University, Uppsala, Sweden; Department of Cell and Molecular Biology, Uppsala University, Uppsala, Sweden; Department of Surgical Sciences, Uppsala University, Uppsala, Sweden; Department of Medical Sciences, Gastroenterology and Hepatology Unit, Uppsala University, Uppsala, Sweden

## Abstract

Interactions between individual pathogenic microbes and host tissues involve fast and dynamic processes that ultimately impact the outcome of infection. Using live-cell microscopy, these dynamics can be visualized to study e.g. microbe motility, binding and invasion of host cells, and intra-host-cell survival. Such methodology typically employs confocal imaging of fluorescent tags in tumor-derived cell line infections on glass. This allows high-definition imaging, but poorly reflects the host tissue’s physiological architecture and may result in artifacts. We developed a method for live-cell imaging of microbial infection dynamics on human adult stem cell-derived intestinal epithelial cell (IEC) layers. These IEC monolayers are grown in alumina membrane chambers, optimized for physiological cell arrangement and fast, but gentle, differential interference contrast (DIC) imaging. This allows sub-second visualization of both microbial and epithelial surface ultrastructure at high resolution without using fluorescent reporters. We employed this technology to probe the behavior of two model pathogens, *Salmonella enterica* Typhimurium (*Salmonella*) and *Giardia intestinalis* (*Giardia*), at the intestinal epithelial surface. Our results reveal pathogen-specific swimming patterns on the epithelium, showing that *Salmonella* adheres to the IEC surface for prolonged periods before host-cell invasion, while *Giardia* uses circular swimming with intermittent attachments to scout for stable adhesion sites. This method even permits tracking of individual *Giardia* flagella, demonstrating that active flagellar beating and attachment to the IEC surface are not mutually exclusive. Thereby, this work describes a powerful, generalizable, and relatively inexpensive approach to study dynamic pathogen interactions with IEC surfaces at high resolution and under near-native conditions.

**Importance:** Knowledge of dynamic niche-specific interactions between single microbes and host cells is essential to understand infectious disease progression. However, advances in this field have been hampered by the inherent conflict between the technical requirements for high resolution live-cell imaging on one hand, and conditions that best mimic physiological infection niche parameters on the other. Towards bridging this divide, we present methodology for differential interference contrast (DIC) imaging of pathogen interactions at the apical surface of enteroid-derived intestinal epithelia, providing both high spatial and temporal resolution. This alleviates the need for fluorescent reporters in live-cell imaging and provides dynamic information about microbe interactions with a non-transformed, confluent, polarized and microvilliated human gut epithelium. Using this methodology, we uncover previously unrecognized stages of *Salmonella* and *Giardia* infection cycles at the epithelial surface.

## Introduction

Although infectious diseases of the intestine are often caused by large populations of invading pathogens, disease progression and outcome are ultimately dictated by the interactions of individual microbes with the host tissues. To characterize the complex dynamics of these underlying interactions, live cell microscopy has become the method of choice. However, it is intrinsically difficult to study dynamic microbe interactions with internal host tissues such as the intestinal epithelium *in vivo*. The resolution of *ex vivo*-based microscopy techniques often suffers from the complexity and depth of intact tissues, resulting in the need for more phototoxic high-dosage illumination or complex and expensive 2-photon setups. Therefore, researchers have turned to transformed or immortalized cell lines to study intestinal epithelial infections in cultured proxies of the gut epithelium. These cell lines often fail to recapitulate key features of intestinal epithelial cell (IECs) layers, such as a densely-packed polarized morphology, a microvilliated apical surface, and sensitivity to cell-death mechanisms, but have nevertheless uncovered a wealth of information about pathogen infection cycles (1–6). By contrast, the impact of physiologically relevant host cell and tissue parameters on infection dynamics remains understudied.

In the past decade, cultured organoid models have been shown to provide a powerful intermediate for this physiological gap between cell lines and intact primary tissues. The central features of the gut epithelium are faithfully recapitulated in both intestinal organoids derived from pluripotent stem cells (PSCs) (7, 8), and in so called enteroids or colonoids derived from adult epithelial stem cells (ASCs) of small intestine or colon, respectively (9, 10). Organoid models can be cultured in a variety of two- and three-dimensional (2D and 3D) settings (11, 12) and retain non-transformed cell behavior over time (13, 14).

Despite their potential, organoid models have so far only been sparsely used for live cell imaging of intestinal infection processes (15–17). In our experience, this is a result of the difficulty to adapt current imaging approaches from cell line infections to accommodate the properties of physiologically grown organoid-derived epithelia. First, while cell lines can be grown flat on glass culture ware for optimal working distance and numerical aperture, intestinal organoid-derived IECs only develop into their natural polarized arrangement when cultured within rich extracellular matrices (ECMs), or atop permeable supports. The latter can be accommodated by ECM-coated transwell inserts with permeable membranes of Polyethylene Terephthalate (PET) or similar polymers as a 2D substrate (18, 19). Enteroid/Colonoid-derived IEC layers in such transwell inserts can be efficiently cultured, differentiated, and infected by a variety of gut pathogens (20–28), but are poorly compatible with live cell imaging. Secondly, non-transformed cells are difficult to manipulate genetically, which complicates introduction of fluorescent tags to visualize the host cell and its subcellular architecture by fluorescence microcopy. Finally, in contrast to tumor-derived cell lines, non-transformed cells retain sensitive cell death and stress signaling pathways, which makes them susceptible to phototoxicity and other perturbations and introduces a need for gentle imaging conditions. Taken together, these constraints have limited the applicability of live cell imaging to characterize encounters of pathogens with the apical portion of non-transformed IEC layers at high spatial and temporal resolution.

Here, we present a new method to visualize microbial infection cycle dynamics at the apical surface of ASC-derived IEC layers under near-native conditions. To overcome the imaging constraints introduced by PET transwells, we developed a method to grow microvilliated human epithelium layers on ECM-coated alumina membranes in 3D-printed imaging chambers. In addition, we omitted the need for fluorescent tagging and high-dosage illumination by optimizing the conditions for high resolution, live differential interference contrast (DIC) microscopy. The use of DIC rather than more phototoxic fluorescent reporter approaches favors physiologic pathogen and host cell behavior and allows simultaneous visualization of both individual microbes and IEC surface ultrastructure without the need for channel switching. Finally, we used this method to map *Salmonella enterica* Typhimurium (*Salmonella*), and *Giardia intestinalis* (*Giardia*) infection cycles atop the epithelial surface and describe previously unrecognized single-microbe behaviors during IEC attachment. Thereby, we show that this imaging methodology enables detailed, dynamic studies of both microbe and host cell behavior at the interface of gut infection, adaptable even to genetically non-tractable microorganisms.

## Results

### An alumina membrane chamber enables high-definition live cell DIC imaging at the apical border of intestinal epithelial cell layers

We developed a method for live cell imaging of microbe interactions with non-transformed human ASC-derived intestinal epithelial cell layers, aiming to combine i) high structural definition, ii) high temporal resolution, iii) gentle imaging conditions to avoid phototoxicity and the need for fluorescent reporters, and iv) a confluent, polarized IEC arrangement. To achieve this, we sought to improve upon the constraints presented by PET transwell supports to DIC imaging (Fig 1A). Specifically, the PET membrane depolarizes light, the pores in the membranes diffract light, and the plastic membrane holder prevents close approach of the microscope objective to the apical side of the IECs, thereby constraining the working distance and numerical aperture. Therefore, a suitable alternative to PET membranes should not introduce optical interference for imaging within the visual spectrum, but like PET membranes should be permeable to permit efficient epithelial cell polarization. In addition, the membrane holder should allow close proximity of both the objective and condenser to the epithelial surface to optimize the numerical aperture of the system.

**Figure 1.**
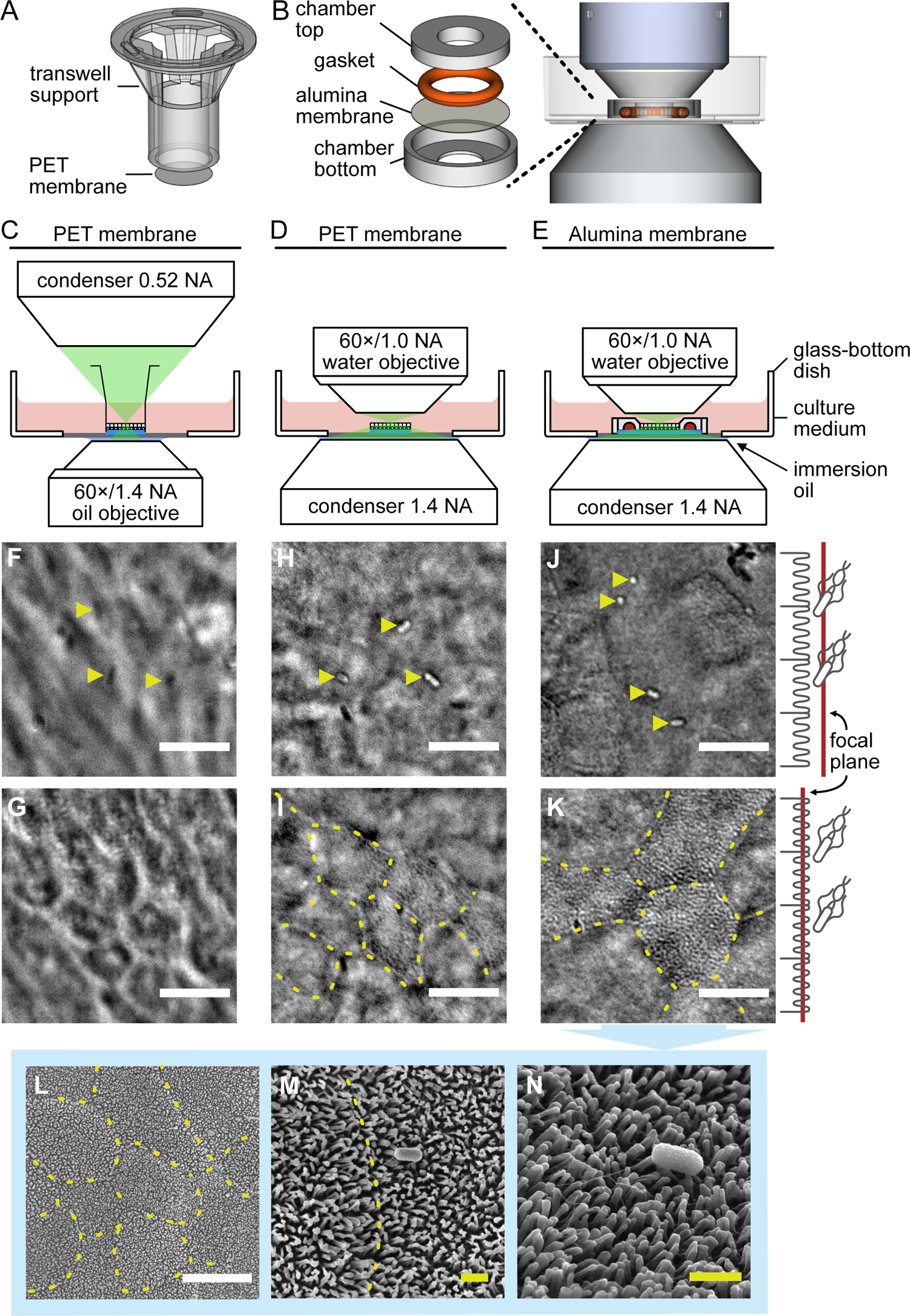
High-definition live cell DIC imaging of IEC monolayer infections in a novel alumina membrane chamber. Schematic comparison of assemblies for IEC monolayer culture highlight that PET transwell membrane holders have a large plastic support (A), which has been omitted in the novel AMCs (B). Both structures were placed in a 35 mm glass bottom dish for imaging. The lower height of AMCs allows for closer proximity, and thus greater numerical aperture, of the microscope’s objective and condenser. The optical interference caused by the large plastic transwell support and resulting larger working distances of both objective and condenser is illustrated by live cell imaging of infections in intact PET transwells in an inverted microscope (C,F,G) versus cut-out PET membranes using an upright water-dipping microscope (D,H,I). This change in microscope setup improved the lateral resolution of bacteria (*Salmonella*) on the cell surface (F,H) and the apical surface topology of IECs (G,I). Either bacteria or surface topology could be emphasized by slightly changing the focal plane, as indicated on the right. Replacing the PET membrane with an alumina membrane held within the custom-designed AMC (E) further improved the imaging resolution and minimized optical interference (J,K). AMCs allow for sequential imaging of the same sample with DIC and SEM (L-N). The latter confirmed that differentiated IECs grown in AMCs exhibit a highly interconnected epithelium with a densely microvilliated surface. Bacteria are indicated with yellow arrow heads. Overlaid dashed lines indicate cell-cell junctions. White and yellow scale bars: 10 and 1 µm, respectively.

This led us to evaluate membranes of anodized aluminium oxide (alumina) as a candidate substrate. Alumina forms a dense honeycomb-like structure with parallel, sub-diffraction limit sized pores (Fig S1A), and is optically transparent when wet. Unlike for PET membranes, the pores cannot be distinguished by light microscopy and the alumina does not depolarize transmitted light (Fig S1B). Although alumina membranes poorly support cell adhesion, various adhesion-enhancing surface modifications have been reported for this material (29–31). We found that surface hydroxylation followed by sequential coating with poly-L-lysine and Matrigel enabled efficient attachment and expansion of human ASC-derived IECs atop the alumina membranes (Fig S1C; see methods for details). Furthermore, IECs grown on coated alumina membranes developed into confluent, highly polarized monolayers, reflecting *in vivo* epithelium architecture (Fig S1D). To hold the alumina membrane in place during cell culture, we designed a 3D-printable plastic chamber (Fig 1B). The design of the chamber can easily be adapted to alternative applications, and design files are freely available for download (will be available upon publication). Like a regular transwell, the chamber creates a cell culture area in the middle, but the height of the chamber was kept slim to match the working distance of a water-dipping objective, thereby removing air-liquid interfaces within the light path. In addition, the chamber allows the objective and especially the condenser to be placed in close proximity to the sample, hence maximizing the utilization of the condenseŕs numerical aperture and thus improving the lateral resolution of DIC imaging.

To test if this alumina membrane chamber (AMC) and upright microscope setup indeed improved the quality of live DIC imaging of epithelial infection, we compared this system to the existing PET transwell supports. Human IEC monolayers were grown and differentiated atop PET transwells, or in AMCs, and the apical compartment was infected with wild-type *Salmonella*. The PET transwells were imaged using either a standard inverted DIC microscope and an oil-immersion objective (Fig 1C), or the PET membrane cut out from the holder for imaging with the upright water-dipping objective setup (Fig 1D). Infections in the AMCs were imaged in parallel using the same upright system (Fig 1E). As expected, the contrast and resolution of both the apical IEC surface and the *Salmonella* was poor for PET transwells imaged through the inverted microscope (Fig 1F-G), but markedly improved through the water-dipping upright system (Fig 1H-I). However, residual optical interference from the PET membrane was still evident, resulting in image blurring. Moreover, the need to cut-out the PET membrane prior to imaging complicated sample handling, increased risk of mechanical cell damage, and caused the loose membrane to float with convection currents in the imaging medium, which prevented stable image acquisition over time.

The AMC markedly improved on all these imaging issues by providing a stable surface, minimizing the optical interference from membrane pores, and allowing easy handling underneath the water-dipping objective. As such, DIC imaging of live, infected IEC monolayers in AMCs showed distinct contrast of *Salmonella* atop the cell surface (Fig 1J). When focusing on the IEC surface itself, we observed clearly demarcated cell-cell junctions and a remarkable roughness made up of contrasting punctae (Fig 1K), suggestive of a densely microvilliated surface. To correlate the live DIC image with the IEC surface topology, we disassembled non-infected and *Salmonella*-infected AMCs and imaged the epithelial monolayers therein also by scanning electron microscopy (SEM). This provided a powerful, near-correlative setup, as SEM analysis could be done on the same AMC samples used for live DIC. SEM images captured at similar magnification validated the appearance of the apical epithelial surface in live DIC mode, both with respect to the macro-topology of cell junctions, the slight height differences between cells, and the distinctly patterned apical surface (compare figure 1K and L). At higher SEM magnification, we observed a richly microvilliated surface (Fig 1M), with preserved binding of *Salmonella* to the host cells (Fig 1N).

In conclusion, the combination of an upright water-dipping objective microscope and the novel AMC provide a system for simultaneous high definition DIC imaging of both microbe- and host cell-features at the apical surface of human intestinal epithelial cell layers. This approach does not require the use of fluorescent dyes or labeled markers, and allows for convenient semi-correlative SEM on samples fixed at the end of live cell imaging.

### Resolving sub second-scale microbial motility patterns along the apical surface of human intestinal epithelium

Pathogenic gut microbes often use flagellar motility to reach and explore the epithelial surface. Motile pathogen behaviors have typically been studied atop artificial surfaces (e.g. plastic or glass) (3, 32), or occasionally under more physiological conditions (e.g. atop tissue explants (33)). However, under the latter conditions the imaging method relies on fluorescent reporters and high-intensity illumination that come at the price of phototoxicity, and may alter the processes under study, or do not permit simultaneous surface structure visualization. The AMC setup presented here should be ideally suited to study authentic pathogen motility patterns at the IEC surface under minimally perturbing conditions. To leverage this possibility, we explored motility of two microbes atop IEC monolayers grown in AMCs: the smaller bacterial pathogen *Salmonella*, and the larger protozoan parasite *Giardia*.

Peritrichous flagella-driven *Salmonella* motility physically constrains stretches of the bacterial swim path atop surfaces – a phenomenon called near-surface swimming (NSS) (3). Using DIC imaging alone, we could successfully follow *Salmonella* NSS along the apical surface of the epithelium with high frame rates (up to ∼30 frames/sec) at modest light intensity, and with simultaneous visualization of apical epithelial topology (Fig 2A, top row panel). Although bacterial NSS could easily be tracked manually based on the DIC images alone, greyscale images cannot be readily thresholded for automated downstream analysis by particle tracking software, as would be the norm for fluorescence imaging (Fig S2A; (34)). We therefore incorporated a squared temporal median (TM^2^) post-processing filter (see methods) to extract bacterial NSS information from the background of the relatively static apical IEC topology. The resulting filtered image stack enabled both segmentation of motile bacteria and automated particle tracking without the use of fluorescent markers (Fig 2A - bottom row panels, movie will be provided upon publication). Automated tracking of *Salmonella* NSS in the TM^2^ filtered stack showed a variety of curved tracks with a mean speed of 34.7 μm/s (Fig 2B, Fig S2C) and mean turning angle of ∼14.68°/15 μm clockwise (Fig S2E), corresponding to its flagella’s counterclockwise spin (35, 36). Reassessment of this pattern by fluorescence imaging resulted in broadly similar tracks (Fig S2A,C), but importantly did not allow simultaneous visualization of epithelial surface topology. These observations validate and extend previous studies of *Salmonella* motility (3, 33, 36–38), by mapping *Salmonella* NSS parameters atop a physiologically arranged epithelial surface and under minimally perturbing conditions.

**Figure 2.**
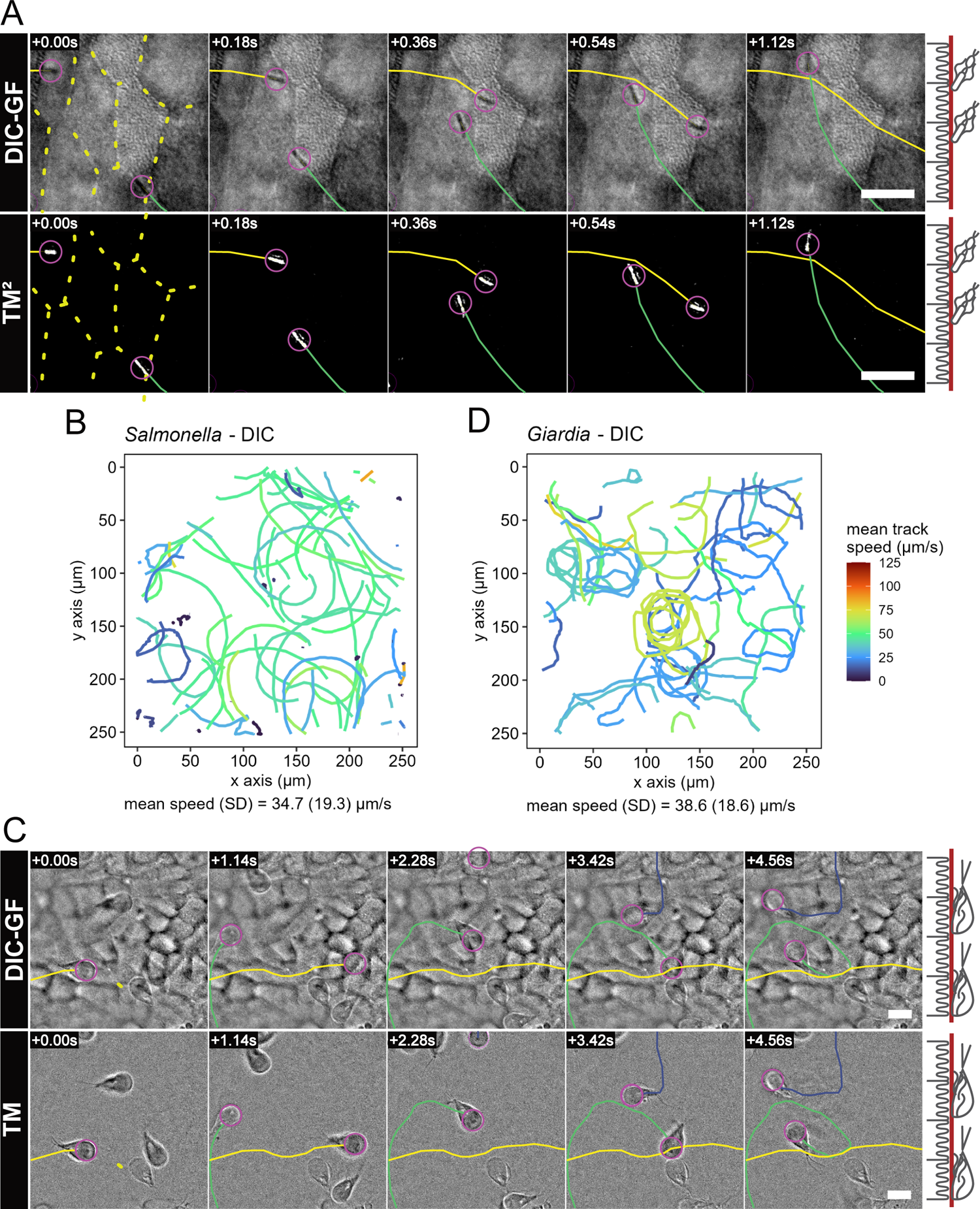
Tracking of microbes on IEC monolayers using DIC imaging resolve *Salmonella* and *Giardia* motility patterns. Confluent, differentiated IEC monolayers in AMCs were infected with *Salmonella-mCherry* (A-B) or *Giardia-mNeonGreen* (C-D). The figure shows tracking from DIC time-lapse movies. Tracking using the respective fluorescent markers (for validation purposes) is shown in Fig S2. DIC movies were processed using a gaussian filter (DIC-GF) to remove uneven background illumination and achieve optimal contrast for *Salmonella* (A, top panel) and *Giardia* (C, top panel). Subsequently, the total DIC-GF stack temporal median projection was subtracted from every frame to specifically emphasize moving structures (TM), and pixel values were squared (TM^2^) to yield positive (white) pixels on a zero (black) background. The TM^2^ filtered time series was used to track all visible *Salmonella* using automated particle tracking (A, bottom panel). For *Giardia*, the TM filtered images were used for manual tracking of swimming trophozoites (C, bottom panel). A random sample of tracks within the field of view (FOV) was used to visualize *Salmonella* motility (B), while all manually tracked paths are shown for *Giardia* (D). The population mean and standard deviation for the track speeds were calculated based on all available tracks (Fig S2C,D). In A and C, a representation of the focal plane (red) is indicated on the right. Tracked microbes are indicated by magenta circles, tracks by continuous colored lines. Overlaid dashed lines indicate cell-cell junctions. Scale bars: 10 µm.

*Giardia* trophozoites feature four pairs of flagella, and use motility to swiftly approach the intestinal epithelium, followed by stable attachment using a ventral disk (39, 40). *Giardia* free-swimming motility in medium involves flexion of the caudal portion of the parasite body (32). On flat surfaces (i.e. glass), *Giardia* adapt the swimming mode to planar motility, largely driven by the flagella. Planar motility in this context is defined by a swimming pattern with the ventral disk continuously in the same plane as the attachment surface (32, 41). As these *Giardia* motility characteristics remain inferred from more simplistic experimental conditions (40), we performed live DIC imaging of early *Giardia* motility atop IEC monolayers within AMCs. Again, we could simultaneously visualize the IEC surface and individual *Giardia* trophozoites, and follow parasite movements at a variety of frame rates (Fig 2C, top row panels). The lower contrast and less predictable swim paths noted for these bigger protozoans, as compared to *Salmonella*, were not well suited for automated particle tracking. However, an unsquared temporal median filtering (TM) step aided robust manual segmentation and tracking of *Giardia* motility (Fig 2C - bottom row panels, movie will be provided upon publication). This revealed epithelium-proximal swimming in curved or circular tracks with a mean speed of 38.6 μm/s (Fig 2D, Fig S2D) and an almost straight mean turning angle of 0.41°/15 μm, but with a large standard deviation in both the clockwise and counterclockwise direction (Fig S2F). We validated these findings also by fluorescence imaging of mNeonGreen-labelled *Giardia* and automated particle tracking (Fig S2B,D). The speed values aligned well with the maximal speeds measured for *Giardia* during free-swimming in media (up to ∼40μm/s; (32)). This indicates that when first approaching a polarized microvilliated epithelium, *Giardia* sustains maximal swim speed for a significant period after switching to planar 2D motility.

Taken together, we demonstrate that DIC imaging in AMCs provides a powerful solution for resolving microbial motility patterns atop a human intestinal epithelial cell layer. Specifically, this methodology simultaneously captures both apical IEC layer topology and single microbe behaviors at high frame rates, does not require fluorescent labelling, and as such avoids both potential problems with reporter toxicity/phototoxicity, as well as the temporal delays that come with switching between multiple imaging channels. Moreover, the AMC imaging technology will allow in-depth analysis of how microbial motility on the epithelial surface is impacted by physiologically relevant surface features (e.g. crevices formed at cell-cell junctions, extruding IECs etc.).

### Longer-term imaging reveals a previously unrecognized Salmonella Typhimurium infection cycle stage atop the epithelial surface

Following NSS, *Salmonella* can adhere to the epithelial surface through a combination of transient interactions via dedicated adhesins and/or flagella, and stable docking to the plasma membrane via type-III-secretion system-1 (TTSS-1) (42–45).

Docking of the TTSS-1 tip subsequently permits the translocation of bacterial effector proteins into the host cell cytoplasm (1). A rich body of work in epithelial cell line models has shown that this leads to the near-instantaneous induction of large actin-dependent membrane ruffles and swift *Salmonella* invasion of the targeted cell (1). However, recent work has also shown that the induced invasion structure phenotype is dependent on the context of the host cell, and suggests a relation between invasion phenotype and host cell polarization status (46). It therefore remains less well understood how the physiological properties of intact non-transformed epithelia may impact *Salmonella* infection cycle stage(s) at the host cell surface.

To survey the longer-term fate(s) of *Salmonella* on IECs, we performed one-hour infections of AMC chamber-grown IEC monolayers. In the resulting movies, we observed an abundance of non-motile bacteria on the surface of the IEC monolayer (Fig 3A). The accumulation of bacteria on the cell surface occurred in a time-dependent manner, and they most often remained attached for the duration of the experiment. On occasion a bacterium was seen to clearly detach, indicating that not all bacteria successfully formed stable docking interactions with the epithelial surface. Surprisingly, stable immobilization of the bacteria only rarely led to successful invasion of the monolayer (Fig 3A). Instead, this analysis uncovered prolonged lingering of *Salmonella* atop the epithelial surface as the predominant behavior, even enabling bacterial division of attached bacteria upon the apical surface (Fig S3). This contrasts sharply to similar experiments in non-polarized epithelial cell line models, where ruffle-dependent *Salmonella* entry begins within seconds to minutes post-attachment (example movie will be provided upon publication) (3, 46). Nevertheless, IEC invasion could still be detected in the AMC-grown IEC monolayers, albeit at lower-than-expected frequency. Some bacteria disappeared from the focal plane without obvious morphological changes to the IEC surface (Fig 3B, top row panels). We could for this category not unequivocally distinguish between sudden detachment or IEC invasion in the absence of overt surface perturbation. In other cases, we observed unambiguous *Salmonella* invasion through a small digitated IEC surface rim transiently formed around the bacterium (Fig 3B - middle row panels, movie will be provided upon publication), or through somewhat more pronounced donut-like ruffles which begun resembling those elicited in *Salmonella*-infected polarized MDCK cells (Fig 3B, bottom row panels) (46–48).

**Figure 3.**
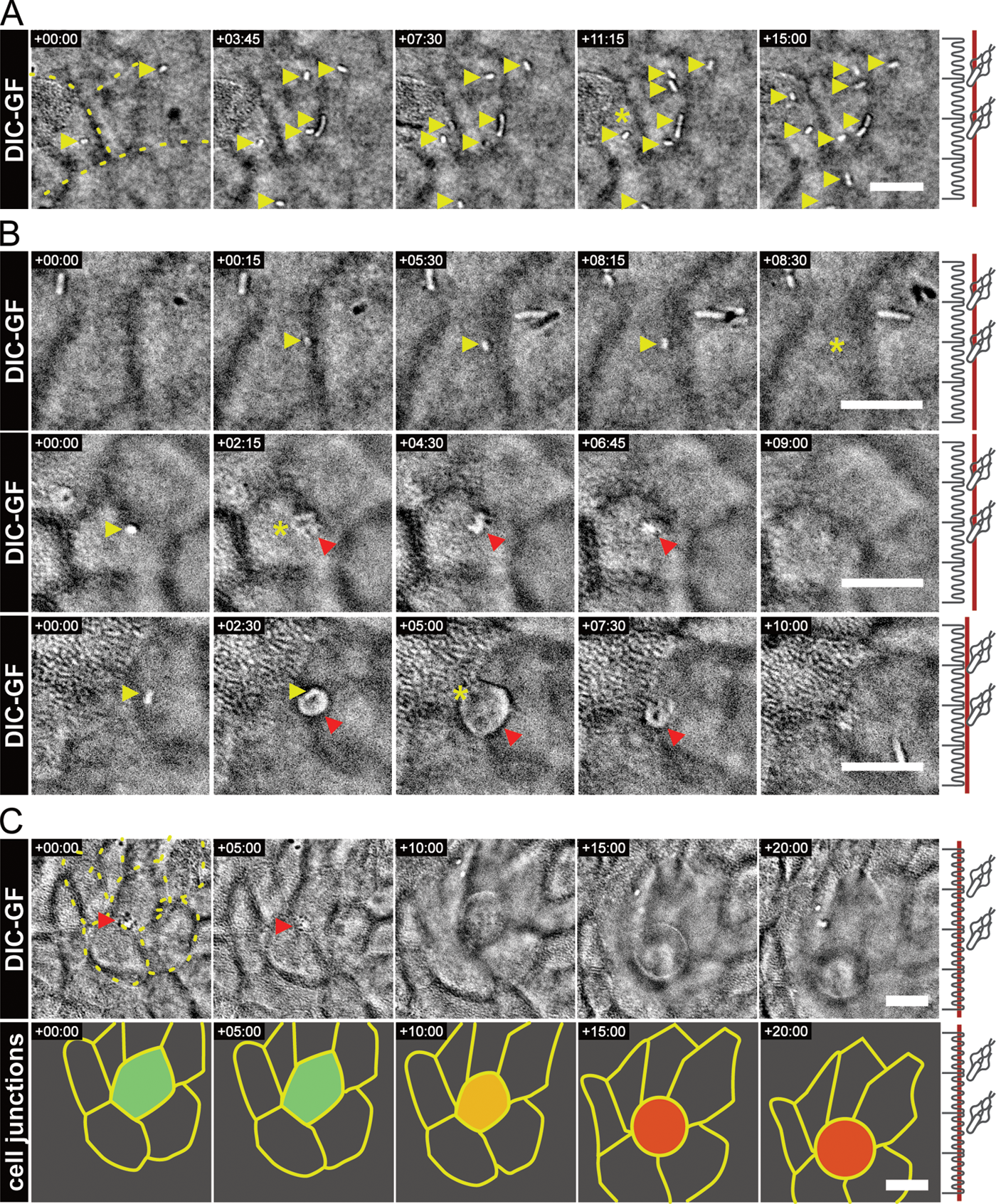
*Salmonella* infection cycle stages at the apical IEC surface. Differentiated IEC monolayers were infected with *Salmonella* and imaged every 15 s. *Salmonella* was most commonly observed to attach to the apical surface for the duration of the experiment (A). A fraction of bacteria was unsuccessful in establishing a lasting adhesion, seen as a short attachment and sudden disappearance (A, asterisk). In other cases, *Salmonella* disappeared from the surface after prolonged attachment either in the absence of visible IEC surface perturbation (B, top row panels), or concomitant with the induction of phenotypically small and discreet (B, middle row panels), or larger donut-shaped (B, bottom row panels) host cell invasion structures. *Salmonella* invasion elicited prompt extrusion of some targeted IECs (C, top panel), which involved an inward movement of the surrounding IECs, evident from an overlaid drawing of the cell-cell junctions (C, bottom panel). Throughout, a representation of the focal plane (red) is indicated on the right. Bacteria are indicated with yellow, and invasion structures with red arrow heads. Yellow asterisks indicate disappearing bacteria (detachment or invasion). Overlaid dashed lines indicate cell-cell junctions. Scale bars: 10 µm.

Successful entry of the bacterium was occasionally followed by prompt neighbor-coordinated extrusion of the targeted IEC from the monolayer (Fig 3C). The extrusion phenotype and timeframe corresponded well with the extrusions we previously reported in *Salmonella*-infected 3D enteroids (15), and which have also been observed *in vivo* (49–51). Again, using DIC imaging alone, the morphological changes in both the extruding IEC and the neighboring cells could be traced over time (Fig 3C).

In conclusion, our results show that imaging of epithelial infection within AMCs opens up new avenues for the microscope-aided study of bacterial behavior at the apical border of a human IEC layer, compatible with both short (sub-second) and long (hours or more) time scales. By employing this technology, we find that *Salmonella* do not invade a physiologically arranged human IEC layer as easily as what has been described for tumor-derived cell line infection models. Instead, stable and prolonged bacterial colonization of the IEC surface constitutes a significant infection cycle stage, which only for a fraction of the bacteria converts into productive IEC invasion. The molecular and physiological basis for these observations constitutes an intriguing area for future research.

### Giardia alternates between rapid swimming and intermittent attachment during local surface exploration

Earlier reports of *Giardia* behavior on glass have described the trophozoites’ pre-attachment swimming pattern as circular movements, largely driven by beating of the anterior and ventral flagella and steered by lateral bending of the caudal region (32, 41). Although the eventual attachment of *Giardia* has been validated in numerous reports (40, 52), and also in epithelial cell line cultures (5, 53), exploratory trophozoite behavior that leads to successful stable attachment has not yet been studied atop an intact epithelial surface.

Therefore, we homed in on individual *Giardia* swim tracks in IEC layer regions that harbored both moving and stably adhered trophozoites (Fig 4A, attached *Giardia* indicated by arrows in top panel). The subset of motile *Giardia* was observed to swim in circular tracks which gradually shifted their position on top of the monolayer (Fig 4B).

**Figure 4.**
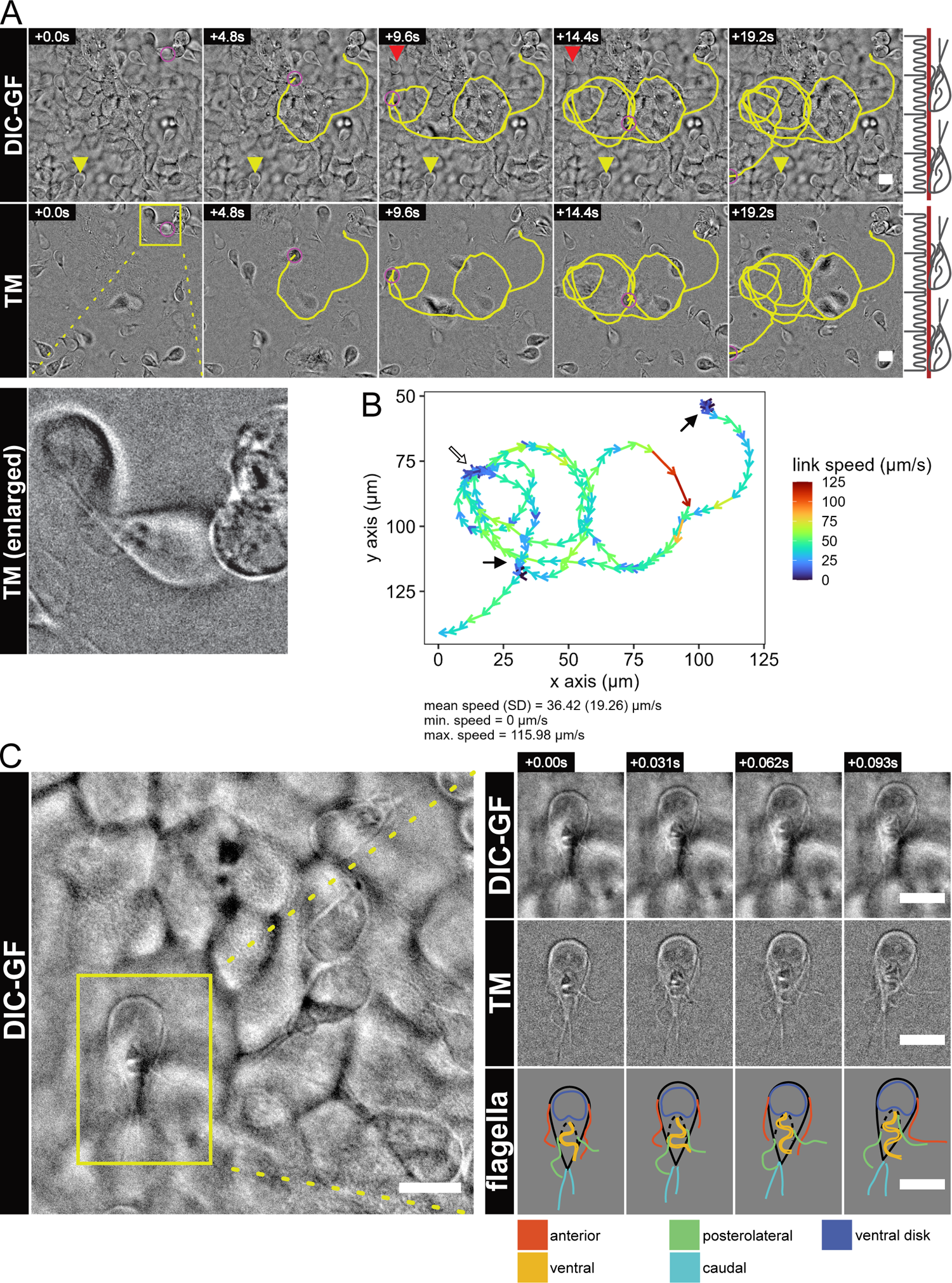
*Giardia* trophozoite exploration of the IEC surface. The link speeds of *Giardia* tracks from Fig 2C,D was analyzed (see also Fig S4) and a representative track showing the common circular pattern was plotted on the DIC (A, top panel) and TM filtered (A, bottom panel and enlarged crop) images. Upon inspection of link speeds, the trophozoite was found to swim with highly variable speeds within a single track (B). The mean, minimum, and maximum speed for this track are indicated (B). During planar swimming, the trophozoite intermittently paused upon sections of the epithelium (B, black arrows), in one location each of the four times the trophozoite visited that particular area (B, white arrow). Upon closer inspection of a single, attached trophozoite (C), the TM filtered time stack showed movement of individual flagellar pairs, which could be manually segmented to follow their shape and position over time (C, middle- and bottom panel to the right, respectively). In A, a representation of the focal plane (red) is indicated on the right. Tracked microbes are indicated by magenta circles, tracks by continuous yellow lines. Yellow- and red arrow heads indicate continuously attached and temporarily attached *Giardia* respectively. The enlarged trophozoite is indicated with a yellow square. Scale bars: 10 µm.

From these tracks, the link speed was calculated as the displacement divided by the time interval between two subsequent frames. Unlike for the patterns described on glass (32, 41), we observed a remarkably high link speed variation for this pre-attachment swimming, with an average of 31.98 µm/s ranging up to ∼155 µm/s (Fig S4). Over time, these tracks scanned repeatedly over a local region of the epithelium (Fig 4A, track).

To study how this swimming behavior relates to IEC attachment, we carefully followed the link speed variation over each full track. This revealed that trophozoites in planar swimming mode occasionally slowed down to a full stop on certain areas of the monolayer (Fig 4B, black arrows). Interestingly, we also found that trophozoites often repeatedly visited a certain location, coming to a brief stop with each pass across that particular stretch of surface (Fig 4B, white arrow). This suggests that some areas of the IEC monolayer may possess properties more favorable to attachment than others. It seems plausible that the highly repetitive circular pre-attachment swim patterns we observe will maximize the parasite’s ability to find such ideal attachment sites within a given epithelial region.

With four pairs of flagella and an adhesive ventral disk, *Giardia* behavior at a surface is the result a complicated interplay of propulsion and adhesive forces. Aside from their role in propulsion, flagellar movement has been described to influence correct positioning of the adhesive disk on the attachment surface, although the exact mechanism of that process remains a topic of discussion (32, 52). To test if our AMC imaging setup would allow studies of individual flagella movements within the context of epithelial infection, we imaged individual *Giardia* atop the IEC monolayer at high frame rate. We found that DIC imaging alone was indeed sufficient to distinguish the movement of individual flagella on intermittently attached trophozoites (Fig 4C, top panel). Application of the TM filter further facilitated manual segmentation of individual flagella (Fig 4C - middle panel, movie will be provided upon publication), and the imaging resolution was sufficient to indicate the movement of all four pairs of flagella over time (Fig 4C, middle and bottom panel). This allowed us to determine that all of the anterior, posterolateral, and ventral flagella exhibit continued movement also in intermittently attached *Giardia* on IEC monolayers. Consequently, intermittent *Giardia* pausing at the IEC surface is not caused by the temporary cessation of flagellar beating. This finding also illustrates the power of the AMC imaging technology to resolve dynamic host-pathogen interactions under near-native conditions even with subcellular resolution.

## Discussion

Recent work has shown that intestinal organoid-derived monolayers grown on permeable supports provide a physiologically relevant culture model for host-pathogen interaction, applicable to both bacteria (21–26, 28), parasites (27), and viruses (20). In this context, organoids bridge the gap between the complexity and polarized nature of the epithelium encountered *in vivo* and the convenient (yet sometimes misrepresentative) properties of continuous cultured cell lines. Furthermore, 2D organoid cultures offer an attractive alternative to 3D culture specifically for studies of microbe interactions with the apical cell surface. The latter can be microinjected (15, 54, 55), fragmented (56), or inverted (57), but because of the 3D-topology these structures are better suited for population-scale dynamics than single-microbe behavior in a stable apical plane. While live-cell imaging of microbe dynamics has been published on monolayers of continuous cell lines (3, 47, 58) and to an extent in microinjected 3D enteroids (15), dynamic behaviors of microbes at the surface of a physiologically arranged human intestinal epithelium remain poorly investigated. We show that this short-coming can be explained by the inherent conflict between the technical prerequisites for high-definition light microscopy (e.g. a short working distance, a thin specimen, and an optically inert substrate) and the current best practices for establishment of polarized IEC layer cultures.

Therefore, we here developed a custom imaging chamber based on a coated alumina membrane substrate. This AMC is compatible with water-dipping objective imaging to maximize optical resolution, while the alumina membrane has optical properties superior to permeable plastic cell culture supports. With these improvements to the ASC-derived IEC monolayer culture system, we present an optimized method for high-definition imaging of host - microbe interactions at the apical surface of a confluent, polarized, non-transformed, human gut epithelium. Finally, we used this technique to uncover previously unknown behaviors of both *Salmonella* and *Giardia* on IEC monolayers.

Our AMC imaging model allows tracing of the behavior of individual *Salmonella* cells over time from the moment they come in proximity to the apical cell surface. Previous work using continuous cell lines and *in vivo* infection models have shown that flagella-dependent *Salmonella* motility drives the approach towards host cells (37, 59) and initiates subsequent NSS-behavior in search of suitable entry sites (3, 33). Our findings regarding *Salmonella*’s curved NSS atop IEC monolayers (Fig 2A,B; Fig S2A,C,E) agree with this model of approach. However, in contrast to observations in e.g. HeLa cells, we found that successful binding to the IEC monolayer does not necessarily lead to prompt ruffle-mediated invasion (Fig 3A). Rather, we observed that the bacteria often remain attached without invading within an hour post-infection. From our current observations, it is not clear if this binding is reversible and adhesin-dependent, or rather comprises irreversible TTSS-1-mediated docking (42). The molecular obstacle(s) that make polarized non-transformed IECs more challenging to invade than typical cell lines constitutes an intriguing area for further enquiry. The prolonged bacterial attachment also suggests an additional biologically relevant epithelial colonization stage in the infection cycle of *Salmonella*, in apparent analogy to attaching-effacing enteropathogens that use the epithelial surface as a primary colonization niche (60).

Nevertheless, we still observed *Salmonella* invasion of IECs (Fig 3B), albeit at lower-than-expected frequency and often through entry structures that were smaller and more transient than previously noted for infections of flat-growing cell lines (example movie will be provided upon publication; (46)). This is in line with the more ‘discreet’ mode of *Salmonella* TTSS-1-dependent IEC invasion described in the intact mouse intestine (46). In agreement with literature (17, 49–51, 61–64), we also observed that successful invasion of the monolayer can lead to immediate extrusion of the infected IEC (Fig 3C). When combined with tools for targeted epithelial cell differentiation, bacterial and mammalian genetics, we expect the AMC imaging technology to provide a framework for deciphering bacterial IEC invasion mechanics and the IEC extrusion response in a near-native epithelial context.

Previous studies of *Giardia* infections have shown that the motile trophozoites can attach to both inert substrates like glass (40) and to microvilliated cell surfaces *in vivo* (65) and in culture (5). Research on interaction dynamics of *Giardia* with inert substrates have shown that flagellar motility is essential for motility, cell division, and selection of suitable attachment sites (32, 41, 52, 66, 67), although this behavior has not yet been studied in real time on live cells. We here report the first study of *Giardia* swimming on top of cultured cells, and on polarized IEC monolayers at that. In general agreement with earlier studies on glass (32), we found that *Giardia* exhibits circular, planar swimming above the attachment surface. Specifically, we found that *Giardia* planar swimming speeds average around 30-40 µm/s with short bursts of up to 155 µm/s (Fig S4). These speeds are much higher than pre-attachment swimming that has been characterized before on glass (32, 41, 67–69). Furthermore, we see circular swimming interceded by short stretches of straight swimming (Fig 4A). Finally, we show that trophozoite planar swimming along the epithelial surface leads to intermittent, transient, attachments to the host cells (Fig 4B). Some of these attachments were seen to repeatedly occur in the same location on the epithelial surface. Therefore, our results suggest that *Giardia* planar swimming is geared towards a local surface-scanning pattern, optimized to select a suitable attachment site within a given region. The circular swimming pattern could aid in this process by repeatedly visiting promising sites of adhesion, until an unknown threshold of stable adhesion is reached. Earlier work on glass has elegantly shown that flagellar beating is instrumental to the planar swimming behavior of *Giardia,* although the respective contribution of the anterior, posterolateral, and ventral pair is still a matter of debate (32, 41). Strikingly, temporal median filtering in our study resolved the continued movements of both anterior, posterolateral, and ventral flagella in intermittently surface-attached *Giardia* (Fig 4C). Although we cannot directly measure attachment force, this suggests that the transient *Giardia* pauses during surface search can be explained by ventral disk or ventrolateral flange (65, 70) engagement with the surface, rather than by an on-off behavior of flagellar propulsion.

As the AMC imaging method can be used to study both the overall swim patterns and the movement of individual flagella, this technique is ideally suited to characterize the complete spectrum of *Giardia* (and other parasite) behaviors atop intestinal epithelia.

Finally, the AMC imaging method can be adapted to accommodate a variety of different host cell types and microbes. As we have shown that the coated alumina membrane supports the growth of ASC-derived human enteroid monolayers, the current system can likely be used to grow a variety of embryonic stem cell-, iPSC- and ASC-derived epithelia. This would allow for high-definition imaging of microbial interactions with any stem cell-derived human epithelium culture (e.g. airway, mammary gland, stomach, or bladder epithelium, see (12) for a comprehensive overview). As ASC-derived organoids retain the locational and functional identity (71) from the donor material, this methodology could also be used to study the impact of congenital disorders of the epithelium (like some forms of Very Early Onset Inflammatory Bowel Disease) on microbial infection. Lastly, recent studies have reported an adapted chamber for ASC-derived intestinal organoid monolayer co-culture with gut anaerobes (26). Innovations like this open the door to even better modeling of both resident and pathogenic microbes under even more physiologically accurate conditions. However, it remains difficult to study dynamic interactions with the host epithelium in this context, as most gut microbiota strains cannot yet be genetically modified to incorporate fluorescent markers. Our AMC approach to image microbe interactions with the intestinal epithelium without fluorescence could here provide a stepping stone for dynamic analysis of host epithelium - microbiota interactions in the coming years.

## Materials & methods

### Ethics statement

Human jejunal ASC-derived enteroids were generated from resected tissue acquired from routine bariatric surgery, following prior informed consent. All personal information was pseudonymized, and the patients’ identities were unknown to researchers working with the tissue samples. These procedures were approved by the local governing body (Etikprövningsmyndigheten, Uppsala, Sweden) under license number 2010-157 with addendums 2010-157-1 (2018-06-13) and 2020-05754 (2020-10-26)

### Alumina Membrane Chambers

The Alumina Membrane Chambers (AMCs) for 13 mm alumina Whatman Anodisc membranes with 0.2 μm pores (GE Healthcare, Little Chalfont, UK) were designed in-house using FreeCAD v0.19 (https://www.freecadweb.org). The chamber comprises a bottom holder and top cover, which fit together using a friction press fit to hold the alumina membrane in place, with culture medium compartments generated on each side of the membrane. The top cover contains a groove to hold a silicone gasket (product number 527-9790, RS Components Ltd. Corby, UK) with inner diameter of 7.65 millimeter in place. AMC design files are available for non-commercial use and will be provided upon publication. The AMC designs were printed using an Original Prusa MINI+ (Prusa Research, Prague, Czech Republic) 3D printer with a 0.4 mm nozzle and a layer height of 0.2 mm. The AMCs were printed in 1.75 mm, clear, natural PLA filament (prod. number 832-0210, RS Components Ltd.).

### Enteroid culture

Human jejunal enteroids were established and cultured as described previously (15, 17). Briefly, pieces of intestinal resections were washed thoroughly in ice-cold PBS and epithelial crypts were subsequently dissociated using Gentle Cell Dissociation Reagent (STEMCELL Technologies, Vancouver, BC, Canada) by nutating at 4°C for 30 minutes followed by trituration. The resulting epithelial fragments were filtered through a 70 μm cell strainer and crypt-enriched fractions suspended in 50 μl Matrigel (Corning, Corning, NY, USA) domes in a 24 well plate. The embedded crypts were cultured in OGM (IntestiCult Organoid Growth Medium (Human), STEMCELL Technologies) with 100 U/ml penicillin/streptomycin (Thermo Fisher (Gibco), Waltham, MA, USA) at 37°C and 5% CO_2_. Growth medium was refreshed every 2-3 days.

For maintenance of human enteroids, the structures were passaged weekly at a ratio of circa 1:8 by mechanical dissociation. The Matrigel domes were manually broken up by pipetting with Gentle Dissociation Reagent and then washed once with DMEM/F12/1.25% BSA. The resulting suspended enteroids were disrupted by triturating 15-20 times with a 200ul pipet tip. Following disruption, enteroid fragments were again suspended in 50 μl Matrigel:Intesticult at the ratio 3:1, divided over 3 domes per well in a 24 well plate, and cultured at 37°C and 5% CO_2_.

### Enteroid-derived IEC monolayer culture

Human IEC monolayers were cultured on either 24-well transparent polyethylene terephthalate (PET) tissue culture inserts with 0.4 µm pores (Sarstedt, Nümbrecht, Germany) or 13 mm diameter alumina Whatman Anodisc membranes. The PET transwell inserts were coated with 40× diluted Matrigel in PBS for one hour at room temperature prior to use. After this time, the coating was completely removed and the cell suspension was immediately added to the transwell inserts. The alumina membranes were coated with extracellular matrix as well, but required more extensive pretreatment. First, the alumina membranes were soaked in 30% H_2_O_2_ for 1 hour at room temperature to add negatively charged hydroxyl groups to the surface (30) and allow protein binding. Then, the alumina membranes were washed in sterile dH_2_O and incubated in 0.1 mg/ml poly-L-lysine (Sigma-Aldrich, Stockholm, Sweden) in dH_2_O for 5 minutes to prepare the surface for Matrigel coating. After poly-L-lysine coating, the membranes were air-dried in a laminar flow cabinet for ∼2 hours or overnight. Finally, the alumina membranes were soaked in 40× diluted Matrigel in dH_2_O for 1 hour, and air-dried again. After coating, the membranes were mounted within AMCs.

Human enteroids were dissociated into single cell suspensions as described before (17). Briefly, circa one well of enteroids per membrane was dissociated into single cells at day 7 after passaging. The enteroids were first taken up from the Matrigel in Gentle Dissociation Reagent, then washed in PBS/1.25% BSA, and dissociated into single cells using TrypLE Express (Thermo Fisher (Gibco)) for 5-10 minutes at 37°C. Cells were then spun down at 300 rcf for 5 minutes and resuspended in OGM+Y (Rho kinase inhibitor Y-27632 (10 μM), Sigma). Finally, the cells were counted manually and 3.0×10^5^ cells where seeded into the apical compartment of PET transwells in 150 μl (600 μl medium in bottom compartment, 24-well plate wells), or into the apical compartment of AMCs in 75 μl (600 μl medium in the bottom compartment, 12 well plate wells). The monolayers typically grew confluent in 2-4 days, whereafter the cells were differentiated towards an enterocyte phenotype by deprivation of Wnt-signalling for 4-5 days. The medium for differentiation consisted of DMEM/F12 supplemented with 5% R-Spondin1 conditioned medium (home made from Cultrex 293T R-spondin1-expressing cells, R&D Systems, MN, USA), 10% Noggin conditioned medium (home made with HEK293-mNoggin-Fc cells, kindly provided by Prof. Hans Clevers, Utrecht University), 50 ng/ml mouse recombinant EGF (Sigma-Aldrich), 1X B27 supplement (Gibco), 1.25 mM N-acetyl cysteine, and 100 U/ml penicillin/streptomycin (9, 10).

### Salmonella Typhimurium strain, plasmid, culture, and infection

All *Salmonella* infections in this study were performed with *Salmonella enterica* serovar Typhimurium, SL1344 (SB300) (72). For validation by standard fluorescence microscopy, the strain carried a pFPV-mCherry (rpsM-mCherry; Addgene plasmid number 20956) plasmid directing constitutive mCherry expression (73). As reported previously, expression of this mCherry construct did not influence motility or invasive behavior of *Salmonella* SL1344. For IEC monolayer infections, *Salmonella* inoculi were grown in LB/0.3 M NaCl (Sigma-Aldrich) for 12 h overnight with 50 µg/ml ampicillin. The following day, a 1:20 dilution was subcultured in LB/0.3 M NaCl without antibiotics for 4h. For subsequent infection of monolayer cultures, the 4h inoculum was diluted to 1.0×10^8^ CFU/ml in DMEM/F12 (Thermo Fisher (Gibco)) without antibiotics of which 10 ul was used for each infection, resulting in 1.0×10^6^ CFU per monolayer.

### Giardia intestinalis culture and infection

*Giardia intestinalis* isolate WB, clone C6 (ATCC 30957) was used in this study. For validation by standard fluorescence microscopy, a modified *Giardia* line constitutively expressing mNeonGreen was generated (see below). *Giardia* trophozoites were grown at 37°C in 10 ml flat plastic tubes (Thermo Fisher Nunc, MA, USA) or 50ml tubes (Sarstedt, Germany) filled with TYDK medium (also known as modified TYI-S-33 or Keister’s medium) (74), supplemented with 10% heat-inactivated bovine serum (Gibco, Thermo Fisher MA, United States). All materials used in the TYDK medium were purchased from Sigma-Aldrich (MO, USA) unless otherwise stated. For IEC monolayer infections, *Giardia* trophozoites were grown until approximately 70% confluence and washed once with TYDK to remove dead cells. Further, trophozoites were incubated on ice (12 min), counted and pelleted by centrifugation (800 x g, 10 min, 4°C). Cells were washed once in 1 ml DMEM/F12 (Thermo Fisher (Gibco)), centrifuged and diluted to 2×10^7^ or 4×10^7^ trophozoites/ml using DMEM/F12 of which 10µl were used for infection, resulting in 2×10^5^ - 4×10^5^ trophozoites per monolayer.

### mNeonGreen plasmid construction and Giardia trophozoite transfection

To visualize *Giardia* trophozoites on IEC monolayers also by fluorescence microscopy, we created a trophozoite strain constitutively expressing mNeonGreen under the control of the beta-giardin promoter. The mNeonGreen gene was PCR-amplified from the pNCS-mNeonGreen plasmid (Allele Biotechnology, CA, USA) and the beta-giardin 5’UTR and 3’UTR regions were amplified from genomic DNA of the WB isolate (see Table S1). The PCR fragments were fused by overlap extension PCR and cloned into the integration vector pPacV-Integ-HA-C (75) using XbaI/PacI restriction sites. *Giardia* trophozoites were electroporated as previously described (76). Transgenic parasites that had the mNeonGreen expression cassette integrated on the chromosome were selected by adding puromycin (50μg/ml) to the culture medium approximately 16 h after transfection. To ensure homogeneous mNeonGreen expression in the culture, we created a clonal trophozoite population from the original mNeonGreen stable transfectant population using serial dilution. Briefly, the original trophozoite culture was diluted in TYDK and seeded as single cells into the wells of a 96-well plate. Wells reaching 70–80% confluence were selected and grown in 10 ml tubes containing TYDK. After the establishment of the clonal mNeonGreen strains the trophozoites were grown without antibiotics.

### Fixed IEC monolayer imaging

Differentiated IEC monolayers grown in AMCs were fixed in 4% PFA for 30 min and permeabilized with 0.1% Triton X-100 for 10 minutes. Then, the cells were stained with phalloidin-AF488 (Thermo Fisher) and 4′,6-Diamidino-2-phenylindole dihydrochloride (DAPI, Sigma-Aldrich) counterstain for 30 min in PBS. Subsequently, the alumina membrane with cells was removed from the plastic membrane holder, washed, and mounted under a 0.17 µm coverslip in Mowiol 4-88 (Sigma-Aldrich). The samples were imaged on a Zeiss LSM700 inverted point-scanning microscope system with a 63X/1.4 NA oil immersion objective using a voxel size of 70.6 nm (x,y) and 0.54 μm (z-sections).

### Live cell infection imaging

Live cell imaging was performed on an inverted Nikon Ti-eclipse microscope (Nikon Corporation, Tokyo, Japan) with a 60X/1.4 NA Nikon PLAN APO objective (0.19 mm WD), Nikon condenser (0.52 NA, LWD), and Andor Zyla sCMOS camera (Abingdon, Oxfordshire, England) with pixel size of 108 nm for Fig 1C,D,F-I and a custom upright microscope for all other experiments. The upright microscope is a custom-build, largely based on the Thorlabs Cerna upright microscopy system (Newton, NJ, USA) with a heated 60X/1.0 NA Nikon CFI APO NIR objective (2.8 mm WD) and a Nikon D-CUO DIC Oil Condenser (1.4 NA) controlled by Micro-Manager 2.0-gamma (77). Images were acquired with an ORCA-Fusion camera (model number C14440-20UP, Hamamatsu photonics, Hamamatsu City, Japan), with a final pixel size of 109 nm. Transmitted light was supplied by a 530 nm Thorlabs LED (M530L3) to minimize phototoxicity and chromatic aberrations. The microscope chamber was maintained at 37°C in a moisturized 5% CO_2_ atmosphere and an objective heater was used additionally. Samples in AMCs were mounted in 35 mm glass-bottom dishes (Cellvis, Mountain View, CA, USA) in 3 ml DMEM/F12 without antibiotics in the microscope’s light path, and allowed to equilibrate for 30 minutes. Then, *Salmonella* or *Giardia* were added in pre-made dilutions directly underneath the objective, and imaging was started immediately with <20 ms exposure times for DIC imaging and <50 ms for fluorescence imaging.

### SEM imaging

For SEM analysis of AMC-grown monolayers, *Salmonella* infected IEC monolayers were fixed at 40 minutes post-infection and compared with uninfected controls. To fix the samples, the monolayers were gently washed once with PBS and fixed at 4°C overnight with 2.5% glutaraldehyde (Sigma) in 0.1M PHEM buffer (60mM PIPES, 25mM HEPES, 10mM EGTA, and 4mM MgSO4·7H20, pH 6.9). Prior to SEM imaging, the samples were dehydrated in series of graded ethanol, critical point dried (Leica EM CPD300) and coated with 5 nm platinum (Quorum Q150T-ES sputter coater). The sample morphology was examined by field-emission scanning electron microscope (FESEM; Carl Zeiss Merlin) using in-lens and in-chamber secondary electron detectors at accelerating voltage of 4 kV and probe current of 100 pA.

### Image processing

The acquired microscopy images were processed with Fĳi (78). DIC images in figures 2, 3, 4, and S3 were filtered to acquire an even field of illumination by subtracting a (30-pixel sigma) gaussian blurred projection from the original. Where indicated, a temporal median (TM) filter was used to extract quickly moving structures (e.g. motile *Salmonella*, *Giardia*, and moving flagella) by subtracting the median projection of the time stack from each individual frame. To enable automated particle tracking of moving *Salmonella* in DIC time series, pixel values of the signed 32-bit TM filtered stacks were squared to convert both ‘dark’ and ‘light’ bacteria in the TM-filtered image to positive values. These TM^2^-filtered images were used for automated particle tracking. Both automated and manual particle tracking was performed with Trackmate for ImageJ v6.0.1 (34).

### Tracking analysis and statistics

Data analysis was performed with R (79) and RStudio (80) and packages available from the Comprehensive R Archive Network (CRAN, https://cran.r-project.org/). Tracking statistics were exported from Trackmate, and actual frame intervals were obtained from the image metadata using the Cellocity python package (81) to correct for variations between frames which would be reflected in inaccurate link speeds. Subsequently, the track data was formatted and plotted in R with the ‘tidyverse’ (82) package. Tracks with a mean speed of >100 µm/s were manually confirmed to be false positives of automated tracking, and therefore excluded from analysis. Track angle changes for DIC-tracked swimming paths were calculated using functions based on the ‘trajr’ package (83) with an interval of three links. For the *Salmonella* swimming tracks, only tracks with a mean speed of >5 µm/s were included in the analysis to filter away most non-motile bacteria. The three-link angle change was divided by the distance travelled from the reference link to calculate the angle change per micron. To aid interpretation of this parameter, the angle change / micron was normalized to angle change / 15 micron, the average distance travelled in 3 links. Other supporting packages for R include ‘ggpubr’, ‘rstatix’, ‘ggbeeswarm’, ‘viridis’, ‘ggridges’, and ‘scales’.

## Acknowledgements

Confocal imaging was performed at the BioVis microscopy core facility of Uppsala University. For SEM imaging at the NMI-UCEM (National Microscopy Infrastructure and Umeå Centre for Electron Microscopy) we express our gratitude towards Cheng Choo Lee for image processing and acquisition, Sara Henriksson for support with the fixation protocol, and Linda Sandblad for advice and facilitating the experiments.

## Contributions

Conceptualization: JMvR, JE, MES; Methodology: JMvR, JE, JG, MES; Investigation: JMvR, JE, JG; Formal analysis: JMvR, JE, JG, MES; Resources: MS, DLW, PMH, SGS, MES; Supervision: SGS, MES; Project administration: MS, DLW, PMH, SGS, MES; Funding acquisition: SGS, PMH, MES; Visualization: JMvR, JG; Writing - Original Draft: JMvR, MES; Writing - Reviewing & Editing: all authors.

