## Supplemental information for "High definition DIC imaging uncovers transient stages of pathogen infection cycles on the surface of human adult stem cell-derived intestinal epithelium"

Affiliations:

*Tables and table legends*

**Table S1. Sequences of the primers used for the construction of the *Giardia*-mNeonGreen line.**

| Primer Name | Sequence (5' to 3') | Description |
|---|---|---|
| Fwd-p-BGiardin-XbaI | ttctctagagtatgcagcactcacagagagatg | Forward primer for Beta-Giardin promoter amplification including XbaI restriction site |
| Rv-p-BGiardin-mNeonGreen | ctcctcgcccttgctcaccatccttttattttcttaacttgggc tcaaat | Reverse primer for Beta-Giardin promoter amplification including mNeonGreen overlap |
| Fwd-mNeonGreen | atggtgagcaagggcgaggag | Forward primer for mNeonGreen cds amplification |
| Rv-mNeonGreen | tacttgtacagctcgtccatgcc | Reverse primer for mNeonGreen cds amplification |
| Fwd-3UTR-BGiardin-mNeonGreen | ggcatggacgagctgtacaagtaagcgcctgcagtaa atcatttac | Forward primer for Beta-Giardin 3'UTR amplification including mNeonGreen overlap |
| Rv-3UTR-BGiardin-PacI | cgtttaattaagtgctcaactgcaaccactac | Reverse primer for Beta-Giardin 3'UTR amplification including PacI restriction site |

*Figure legends*

**Figure S1. Characterization of human IEC monolayers grown within AMCs (supplement to Fig 1).** The alumina membranes that were used feature densely packed pores with a diameter of ~0.2 µm, as shown by SEM (scale bar: 1 µm) (A). A major advantage of the alumina membranes versus the standard PET membranes is the lack of interference with polarized light. As shown in B, PET membranes affect the polarization of transmitted DIC light compared to the polarized light behavior through glass. Alumina membranes do not interfere with the light's polarity, and are in this characteristic indistinguishable from glass. Although alumina membranes do not intrinsically support IEC monolayer culture, surface treatment with $H_2O_2$ and poly-L-lysine (PLL), and a subsequent protein coat of Matrigel (MG) enable attachment of organoid-derived IEC's (C). When enteroid-derived IECs are grown and differentiated on coated alumina membranes, a confluent, polarized, and microvilliated IEC monolayer is formed (D). The z-stack in C shows the microvilliated surface (red frame), cell-cell junctions (yellow frame), super-nucleus height (green frame), equally distributed nuclei in a continuous plane (cyan frame), and basal stress fibers (blue frame). Nuclei are stained with DAPI, F-actin with phalloidin-AF488. Scale bars: 10 µm.

**Figure S2. A summary of tracked microbe dynamics (supplement to Fig 2).** Confluent, differentiated IEC monolayers were infected with *Salmonella-mCherry* or *Giardia-mNeonGreen*. Time series were acquired in the respective fluorescent channels, and a random sample of *Salmonella* tracks (A) and all *Giardia* tracks (B) within the field of view were used to visualize microbe motility. The population mean and standard deviation for the track speeds were calculated based on all available tracks (C,D). Imaging of the DIC channel for both microbes was used to calculate the turning angle over 15 µm (E,F). To remove angle change measurements from non-motile *Salmonella*, only tracks with a mean track speed of > 5 µm/s were included in the analysis. Since manual tracking of *Giardia* was only performed for motile trophozoites, these tracks were analyzed without filter. The distribution of mean angle changes for *Salmonella* swimming tracks show a central tendency of the bacteria to continue a clockwise (+) rather than counterclockwise (-) circular pattern, with small standard deviation which allows for more ellipsoid tracks and lateral movement (E). For *Giardia*, the angle change distribution does not show a net clockwise or counter-clockwise tendency, but rather an almost straight mean trajectory with large standard deviation towards both directions (F).

**Figure S3. Attached *Salmonella* dividing on the IEC surface (supplement to Fig 3).** Under the conditions reported in Fig 3, attached bacteria lingered on the apical surface of IEC monolayers for long periods without the induction of entry structures. These attached bacteria could even be seen to grow and divide, where mother and daughter cells dissociated and remained attached to the IEC surface throughout the division cycle. Images are shown in both the DIC and mCherry channel. Scale bars: 10 µm.

**Figure S4. Summary of the link speeds of manually tracked *Giardia* (Supplement to Fig 4).** The individual, manually tracked paths of *Giardia* trophozoites from the DIC imaging shown in Fig 2D were used to analyze the relative frequency of link speeds.

Shown are the relative frequency distributions (kernel density estimation) of link speeds, ordered by the mean speed for each individual track. Most tracks show more than one maximum, indicating that trophozoite movement occurs at different speeds for various stretches within one track. The shown population mean, minimum, and maximum speed were calculated on the collection of all links, regardless of their parent track.

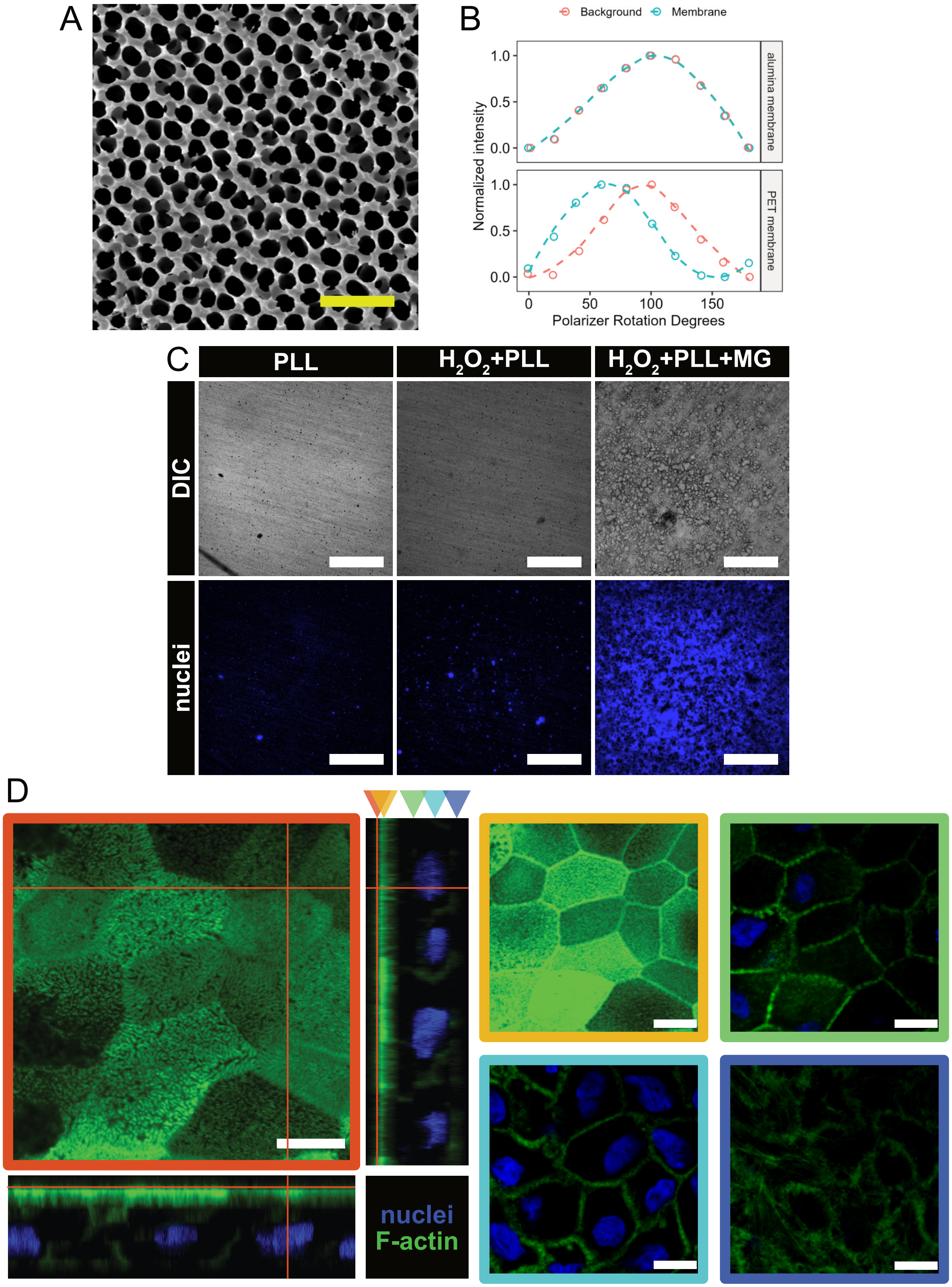


**Figure S1. Characterization of human IEC monolayers grown within AMCs (supplement to Fig 1).**

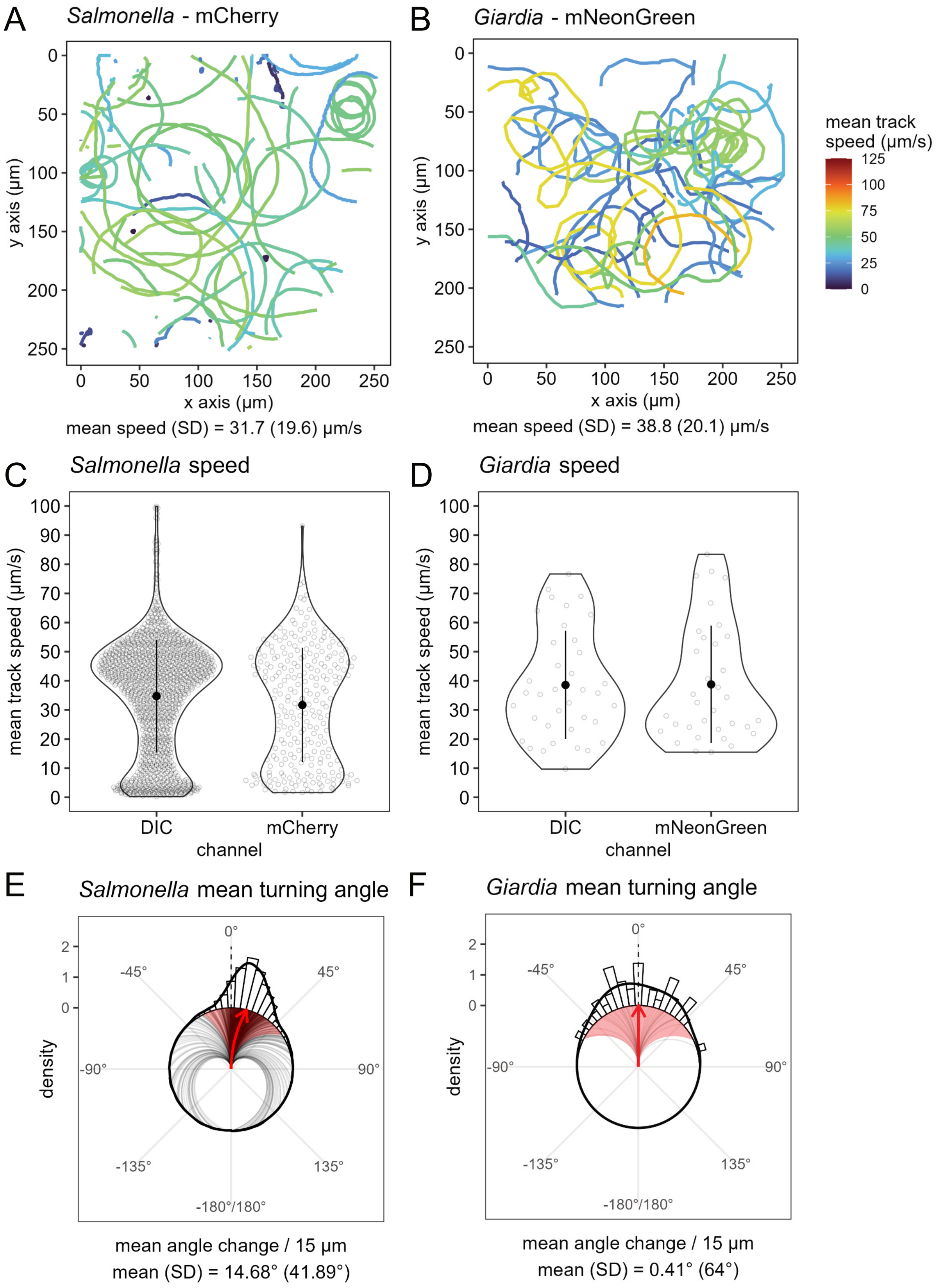


**Figure S2. A summary of tracked microbe dynamics (supplement to Fig 2).**

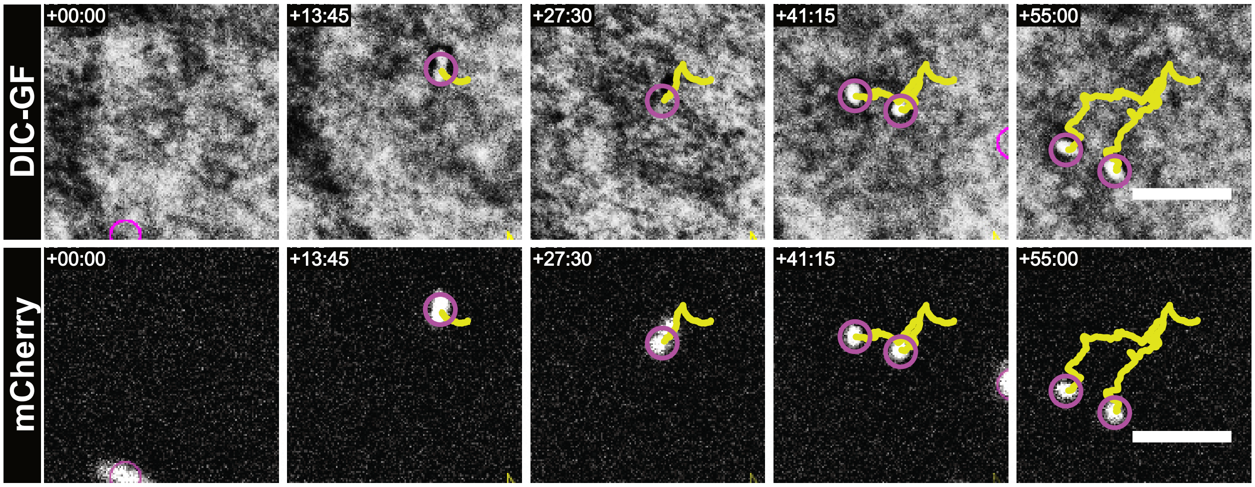


**Figure S3. Attached Salmonella dividing on the IEC surface (supplement to Fig 3).**

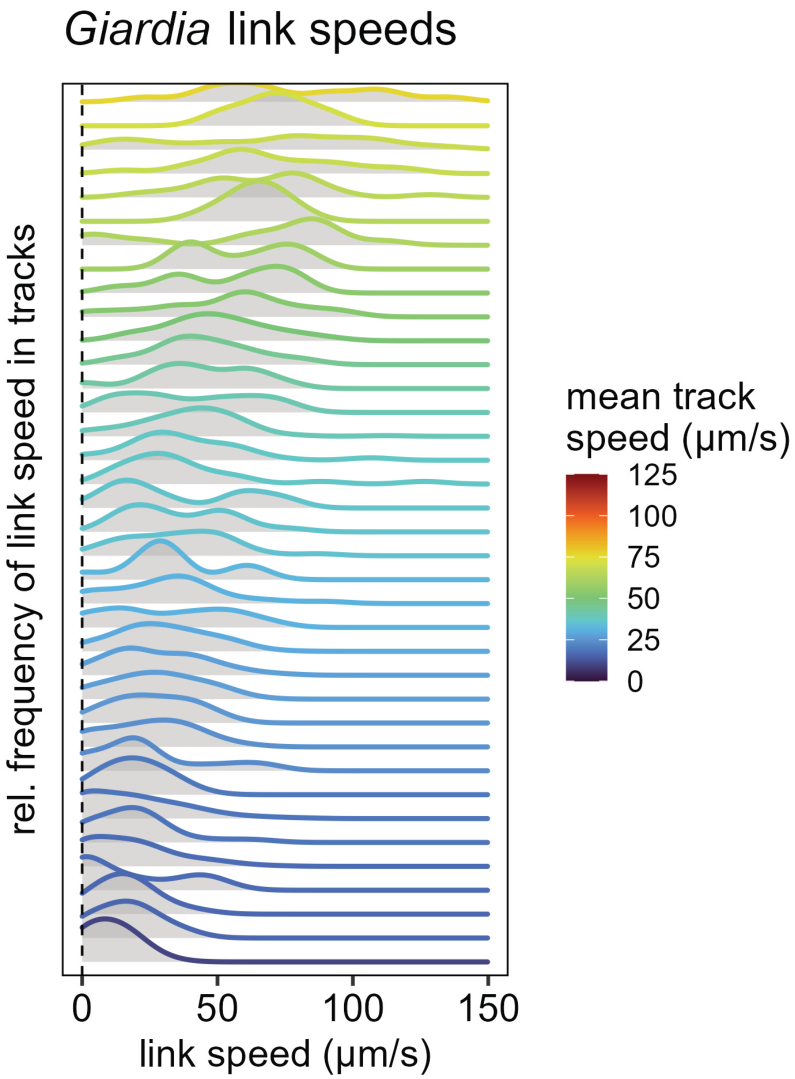


pop. mean speed = 31.98 µm/s
pop. min. speed = 0 µm/s
pop. max. speed = 154.72 µm/s

**Figure S4. Summary of the link speeds of manually tracked Giardia (Supplement to Fig 4).**

*Supplemental Methods*

**IEC attachment assay**

IEC monolayers were grown as described in the main Methods section, on alumina membranes coated with PLL alone, $H_2O_2$ and PLL, or $H_2O_2$ + PLL + MG. After three days of growth in OGM+Y the monolayers were fixed in 4% paraformaldehyde and stained for nuclei with DAPI. Imaging was performed for DIC and DAPI fluorescence on a custom-built microscope based on an Eclipse Ti2 body (Nikon), using 4X/0.2 NA Plan Apo Lambda air objective (Nikon), and a back-lit sCMOS camera with a pixel size of 2.8 μm (Prime 95B, Photometrics). Fluorescence was excited using a Spectra-X light engine (Lumencor) and emission was collected through a quadruple bandpass filter (89402, Chroma).

**HeLa cell infection**

HeLa CCL-2 cells (ATCC) were maintained in DMEM with 10% FCS and 100 U/ml penicillin/streptomycin in plastic culture flasks and passaged every 2-3 days. For infection, $2.0\times10^4$ cells were seeded on 13 mm alumina Whatman Anodisc membranes with 0.2 μm pores in AMCs. Infection was performed the next day with $1.0\times10^6$ CFU *Salmonella* in DMEM. Imaging was performed in the upright water-dipping microscope as reported in the main materials and methods section.
